# Theory of cytoskeletal reorganization during crosslinker-mediated mitotic spindle assembly

**DOI:** 10.1101/419135

**Authors:** A. R. Lamson, C. J. Edelmaier, M. A. Glaser, M. D. Betterton

## Abstract

Cells grow, move, and respond to outside stimuli by large-scale cytoskeletal reorganization. A prototypical example of cytoskeletal remodeling is mitotic spindle assembly, during which microtubules nucleate, undergo dynamic instability, bundle, and organize into a bipolar spindle. Key mechanisms of this process include regulated filament polymerization, crosslinking, and motor-protein activity. Remarkably, using passive crosslinkers, fission yeast can assemble a bipolar spindle in the absence of motor proteins. We develop a torque-balance model that describes this reorganization due to dynamic microtubule bundles, spindle-pole bodies, the nuclear envelope, and passive crosslinkers to predict spindle-assembly dynamics. We compare these results to those obtained with kinetic Monte Carlo-Brownian dynamics simulations, which include crosslinker-binding kinetics and other stochastic effects. Our results show that rapid crosslinker reorganization to microtubule overlaps facilitates crosslinker-driven spindle assembly, a testable prediction for future experiments. Combining these two modeling techniques, we illustrate a general method for studying cytoskeletal network reorganization.

## Introduction

Cell survival depends on cells’ ability to divide, move, grow, and respond to changing conditions, biological functions enabled by flexible and rapid remodeling of the cellular cytoskeleton. Cytoskeletal remodeling is essential for polarized growth, both of single cells [1, 2] and tissues [3]. During cell crawling and adhesion, turnover of the actin and microtubule cytoskeletons is tuned by signaling events [4, 5]. Large-scale cellular volume changes required for phagocytosis, exocytosis, and endocytosis require actin remodeling [6]. Proper organization and localization of organelles, including mitochondria [7, 8] and the endoplasmic reticulum [9], depend on dynamic interactions with the cytoskeleton. During mitosis and cytokinesis, the cytoskeleton undergoes large rearrangements to construct the mitotic spindle [10] and cytokinetic ring [6]. Given the ubiquity of cytoskeletal remodeling for cellular behavior, it is not surprising that aberrant cytoskeletal dynamics or regulation are associated with many diseases, including cancer and developmental defects. The ability of the cytoskeleton to undergo rapid and large structural rearrangements is therefore of broad importance in biology.

The dynamic reorganization of cytoskeletal assemblies is facilitated by a small number of building blocks—filaments, molecular motors, crosslinkers, and associated proteins—which work in concert to generate force [4]. Cytoskeletal filaments, including microtubules (MTs) and actin, undergo dynamic cycles of polymerization and depolymerization that allow rapid turnover. Filament nucleation is controlled by cellular factors (such as *γ*-tubulin for MTs [10] and the Arp2/3 complex for actin [6]), which spatiotemporally tune filament localization. Crosslinking proteins, including diverse actin crosslinkers [5, 11] and the Ase1/PRC1/MAP65 family of MT crosslinkers [12–18], bundle cytoskeletal filaments to organize higher-order assemblies. These proteins can also have a preferred polarity for crosslinking parallel or antiparallel filament pairs [5, 13]. Motor proteins such as kinesins and myosins can walk on filaments to transport cargo, or crosslink and slide filaments to reorganize them [4]. MT-and actin-binding proteins often regulate filament length and dynamics, allowing further control of cytoskeletal architecture [19–23]. While the flexible, dynamic nature of the cytoskeleton is essential for its biological function, we currently lack predictive understanding of cytoskeletal reorganization.

Biophysical modeling of cytoskeletal systems can reveal molecular mechanisms essential for self-assembly, give insight into the physical requirements for a given behavior, and can be dissected in more detail than is possible in experiments. For example, mathematical modeling has been used to study traveling waves of actin polymerization and protrusions in cell motility [24, 25], cortical MT organization in plants [26–29], mitosis [30, 31] and cytokinesis [32, 33]. A frontier in cytoskeletal modeling is the development of general methods to handle three-dimensional systems in which large structural rearrangements occur. Early mathematical modeling of the cytoskeleton focused on one-dimensional problems such as muscle contraction [34–36] and mitotic-spindle length regulation [37–45]. However, higher-dimensional models are required to capture significant rearrangements, such as the actin protrusions of motile cells, cytokinetic ring formation, and cell cleavage.

A prototypical example of cytoskeletal reorganization is mitotic spindle assembly, during which spindle MTs reorganize to form a bipolar structure as the spindle poles separate (Figure 1) [46]. This requires significant structural rearrangement, even in yeasts, which enter mitosis with side-by-side spindle-pole bodies (SPBs), the nucleating centers for spindle MTs (Figure 1B) [47, 48]. In the closed mitosis of yeasts, the SPBs are embedded in the nuclear envelope and therefore undergo constrained motion [49]. Steric interactions between MTs, SPBs, and the nuclear envelope, along with fluid drag from the nucleoplasm, resist large-scale rearrangement of spindle components [50, 51]. Motor proteins and crosslinkers that bundle and slide MTs are key force generators for spindle assembly that create, extend, and stabilize MT bundles [13, 17, 40, 47, 48, 50, 52–62].

**Figure 1:**
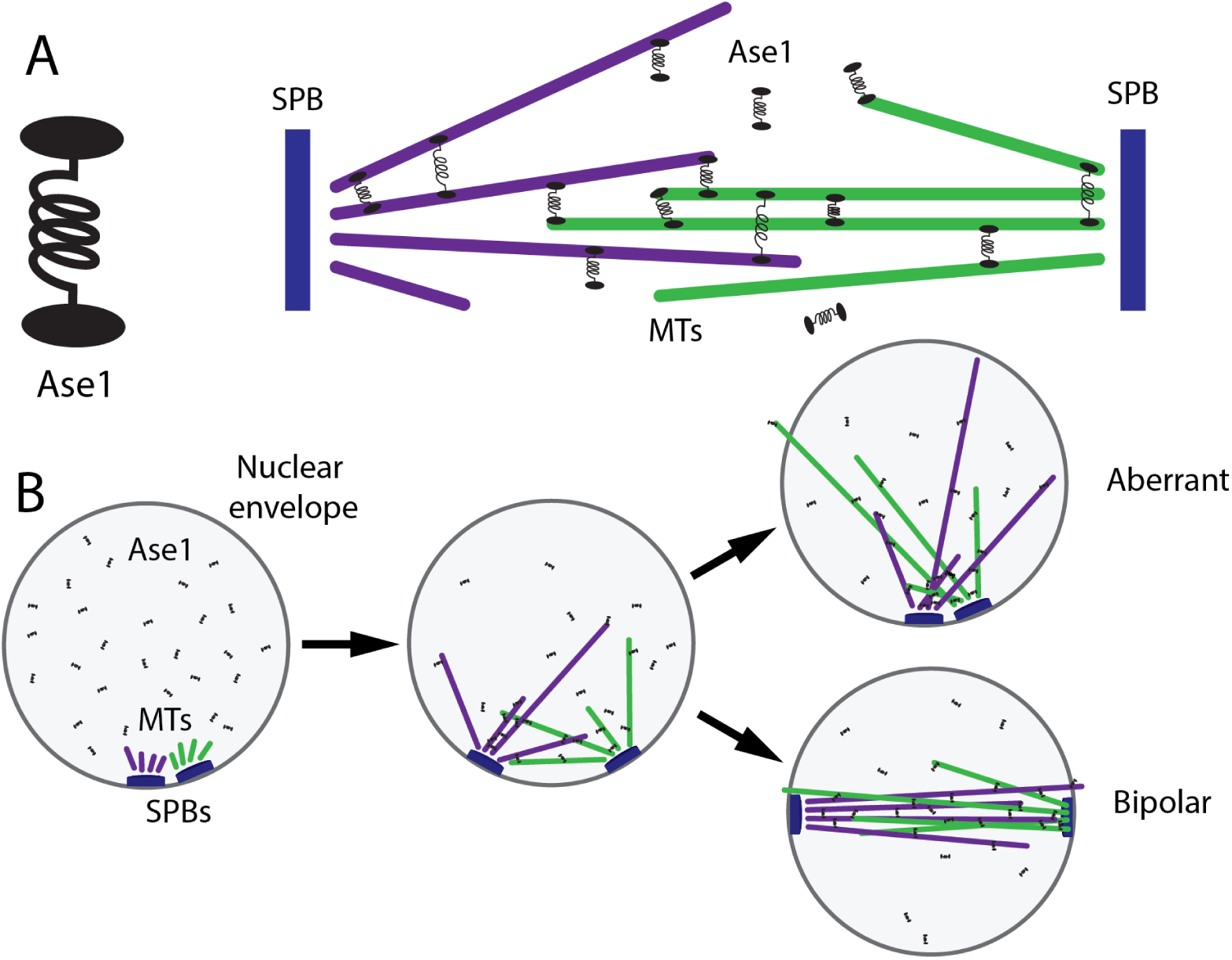
Schematic of spindle assembly by passive crosslinkers. A, Ase1 crosslinker (left) and mitotic spindle (right), including spindle-pole bodies (SPBs, blue), MTs (purple and green), and Ase1 (black). B, Steps of spindle assembly starting from side-by-side SPBs (left), which leads to either a kinetically trapped aberrant state (right, top) or a bipolar spindle (right, bottom)

In contrast to the previously established importance of molecular motors in spindle assembly, recent work showed that fission-yeast spindles can assemble in the absence of mitotic motors if the passive crosslinking protein Ase1 is present [51, 63]. In most organisms, kinesin-5 motors that crosslink antiparallel MTs and slide them apart are essential for spindle-pole separation and the establishment of a bipolar spindle [47, 48, 52, 64–70]. Therefore, it was surprising that in fission yeast, kinesin-5 deletion mutants become viable when the kinesin-14s are simultaneously deleted, spindle assembly is rescued [53, 71, 72]. In fact, spindle assembly can occur with all mitotic motors deleted or inhibited, and Ase1 is essential for spindle assembly in this context [51, 63]. In previous work, we developed a simulation model of spindle assembly both with [50] and without [51] motor proteins. In these simulations, antiparallel crosslinking by Ase1 and the stabilization of crosslinked MT dynamics are sufficient for model bipolar spindle assembly in the absence of motors [51]. Related previous theory has studied mitotic MT bundling by motors and crosslinkers [73, 74].

Here we build on previous work to interrogate the requirements for crosslinker-mediated spindle as-sembly and characterize how varying properties of crosslinkers change spindle morphology. In addition, we develop new theory to understand the minimal requirements for spindle structural reorganization and bipolar spindle assembly and make quantitative predictions for experiments. In order to describe the requirements for passive crosslinkers to align spindle MTs into a bipolar state, we developed a minimal model that accounts for rotation generated by MT polymerization, crosslinkers, drag, and steric interactions. We find that spindle assembly is sensitive to the initial angle at which MT bundles crosslink: smaller initial angle and increased SPB separation facilitates spindle bipolarity. This leads to a geometric explanation of why spindles do not require extensile forces from motor proteins during assembly.

The model we develop includes the general features of dynamic cytoskeletal filaments, forces and torques generated by crosslinkers, friction, and steric interactions. Therefore, this torque-balance model can be applied generally to model crosslinked filament networks driven by polymerization dynamics, contraction, extension, and filament alignment.

## Methods

### Minimal model of microtubule reorientation during spindle assembly

The forces and torques that drive cytoskeletal reorganization can be modeled and simulated using statistical mechanics and hydrodynamic drag. To create a predictive, tractable model of spindle assembly, we considered forces from interactions of MT bundles, SPBs, the nuclear envelope, and the nucleoplasm (Figures 1, 2, S1). MTs are nucleated by SPBs and subsequently bundled by passive crosslinkers. While in principle multiple bundles may be nucleated from one SPB, there is typically one dominant bundle [75, 76]. SPBs are embedded in the nuclear envelope, which is roughly spherical and confines the SPBs to a position on that spherical shell. Motion of the SPB in the shell encounters a viscous drag (Figures 2, S1). The shell prevents MT bundles and other elements of the nucleoplasm from exiting the nucleus. Nucleoplasmic viscosity produces a drag force that opposes the lateral motion of MT bundles, and controls crosslinker diffusion (Figure 2A).

**Figure 2:**
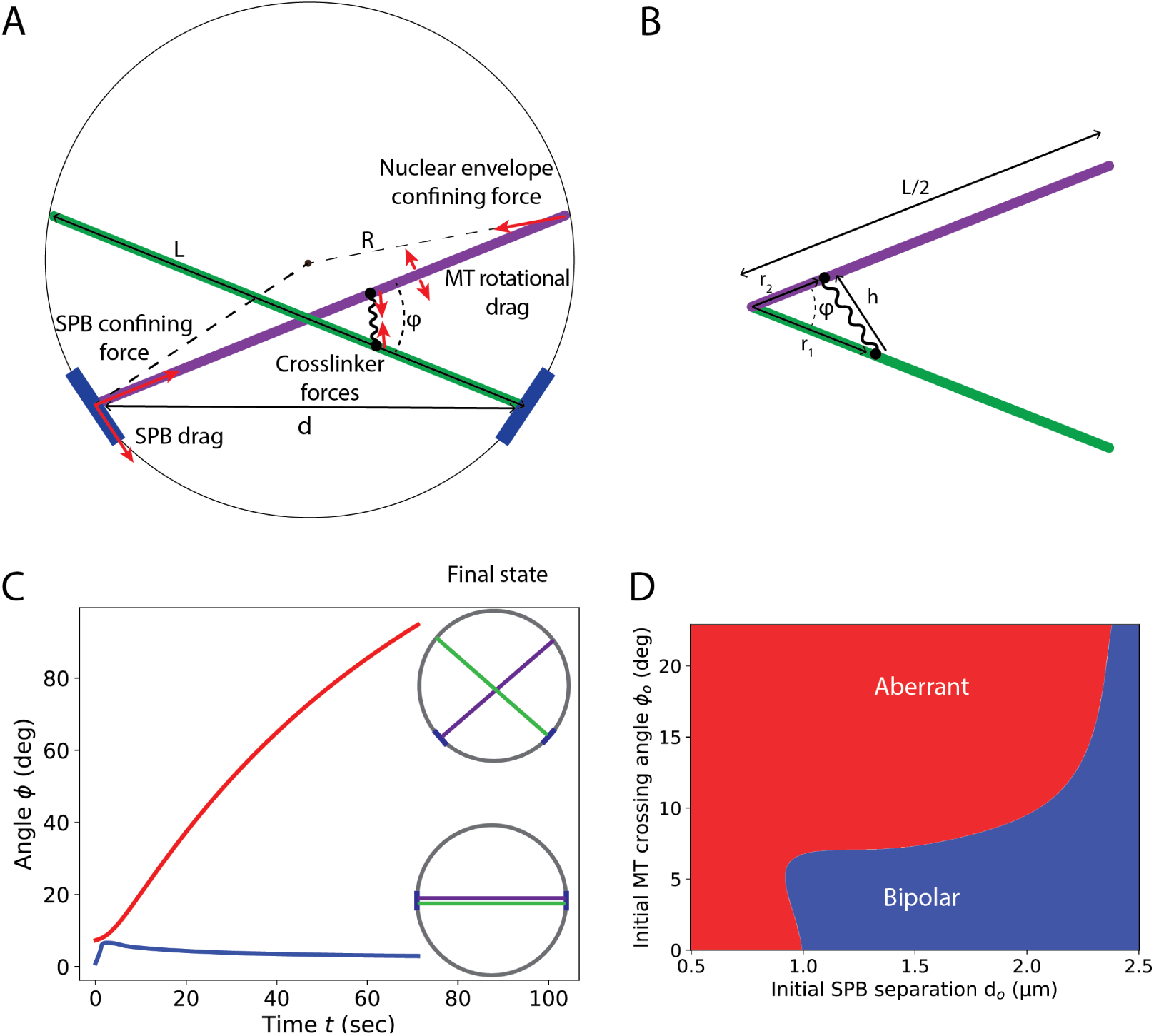
Schematic of and results from the torque-balance model. A, Schematic of forces and torques. MT bundles of length *L* are attached to SPBs, separated by distance *d* and crossing angle *φ*. Forces arise from crosslinkers, drag, and confinement within the nuclear envelope. B, Schematic of crosslinking geometry. Two MT bundles of length *L* cross at angle *φ*, with crosslinker heads binding at distance *r*_1_ and *r*_2_ from the MT bundle ends. C, MT bundle crosslinking angle *φ* as a function of time in the torque-balance model for two initial conditions. Red (top) larger initial crossing angle becomes larger, leading to an aberrant final state. Blue (lower) smaller initial crossing angle decreases, leading to an aligned bipolar spindle. D, Phase diagram of bipolar spindle assembly in the torque-balance model as a function of initial SPB separation and crossing angle. Blue, bipolar spindles form. Red, aberrant spindles form. Parameters are *v*_p_ = 4 *µ*m*/*min, *h*_cl_ = 53 nm, *k*_cl_ = 0.2055 pN/nm, *R*_NE_ = 1.375 nm, *c*_cl_ = 2.88 × 10^-4^ nm^-2^, *z* = 1, *d*_*o*_ = 0.55 *µ*m, *D*_spb_ = 4.5 × 10^-4^ *µ*m^2^*/*sec, initial angle *φ*_*o*_ = 6.07 degrees for bipolar final state in panel C, 9.1 degrees for aberrant final state in panel C. See table 1.

As an MT bundle grows, steric interactions with the nuclear envelope cause the bundle end to slide along the edge of the nucleus, which moves the bundle. We assume that the longest MT within the bundle touches the nuclear envelope, and that MT ends slide freely along the surface of the nuclear envelope, neglecting friction in sliding studied previously [77–79].

**Table 1:** Table of parameter values used in the torque-balance model. Values were chosen to match experimental measurements and analogous parameters in kMC-BD model.

| Parameter | Symbol | Value | Notes |
| --- | --- | --- | --- |
| Crosslinker spring constant | $k_{cl}$ | 0.2 pN nm<sup>-1</sup> | [17] |
| Crosslinker length | $h_{cl}$ | 53 nm | [17] |
| Crosslinker fugacity | $z_{cl}$ | 1.0 | Chosen to have ~20 crosslinkers per MT at full overlap |
| Crosslinker binding affinity | $c_{cl}$ | $3 \times 10^{-4}$ nm<sup>-2</sup> | Chosen to have ~20 crosslinkers per MT at full overlap |
| Nuclear envelope radius | $R_{NE}$ | 1.375 μm | [75] |
| Nucleoplasm viscosity | $\eta_{nu}$ | 10<sup>3</sup> cP | [50, 75] Supporting material |
| SPB diffusion coefficient | $D_{spb}$ | $4 \times 10^{-4}$ μm<sup>2</sup> sec<sup>-1</sup> | [50] |
| MT stall force | $F_{stall,MT}$ | 14.8 pN | [87] |
| MT polymerization rate | $v_o$ | 4 μm min<sup>-1</sup> | Range 0.5-10.0 μm min<sup>-1</sup> [75, 76] |
| Time step | $\delta t$ | 0.0358 sec | Chosen for numerical stability |

Individual MTs undergo dynamic instability characterized by a polymerization speed *v*_p_, depolymerization speed *v*_d_, catastrophe frequency *f*_*c*_, and rescue frequency *f*_*r*_. In the bounded growth regime, these dynamic instability parameters lead to a decaying exponential distribution of MT length [80], which has been observed for some populations of spindle MTs [81]. Consistent with this expectation, the measured dynamic instability parameters of single fission-yeast mitotic microtubules were in the bounded growth regime [76], and therefore we used these parameters to model dynamics of single MTs in the full stochastic model (Table S1). However, crosslinked spindle MTs can be significantly stabilized, for example by CLASP [57, 82]. As a result, bundled MT dynamics parameters may shift into the unbounded growth regime. Consistent with this hypothesis, the length distribution of midzone and kinetochore MTs in the spindle is nonexponential in several organisms [81, 83–86]. Because unbounded growth may occur in spatially confined regions such as the spindle midzone, we consider dynamic instability parameters in this regime for bundled MTs (Table S1). In the minimal torque-balance model of MT bundles, we neglect single-MT catastrophe and consider an average bundle growth speed *v*_pavg_ of

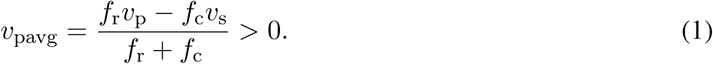

MT polymerization speed slows in response to compressive force along the MT axis, which we model as in previous work (Equation S7) [87]

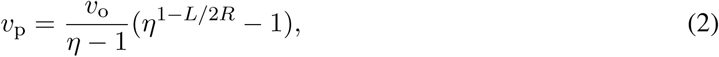

where *v*_o_ = *v*_p_(*F*w, ‖ = 0) is the zero-force polymerization speed and the tubulin association constant *η* = *k*_2,on_*/k*_2,off_ = exp(*δF*_s_*/k*_B_*TN*) where *δ* is the tubulin dimer length of 8 nm, *F*_s_ is the force at which MT polymerization stalls, *N* = 13 is the number of protofilaments in an MT, *k*_B_ is the Boltzmann constant, and *T* is the temperature.

We assume perfectly rigid MTs, neglecting MT and bundle bending. The critical buckling force *F*_c_ = *EI/L*^2^, where *EI* is the flexual rigidity of an MT and *L* its length. In our model, this force arises from the component of the wall force along the MT axis, *F*_w, ‖_ = *F*_*s*_*L/*2*R*, giving a critical buckling length for an MT of *L*_c_ = (2*REI/F*_*s*_)^1^*/*^3^. The flexural rigidity *EI* = 7.9 pN *µ*m^2^ [88], and the nuclear diameter 2*R* = 2.75 *µ*m. If we consider the upper limit of the wall force as the MT growth stall force *F*_*s*_ = 14.8pN, the critical length would be 1.14 *µ*m, approximately the average length of unstabilized microtubules [76]. In practice, force-induced catastrophe of MTs causes MTs to transition to shrinking before reaching the buckling force. Further, in a multi-MT bundle the buckling force greatly increases: for a hexagonal bundle of 14 MTs each 2.75 *µ*m long, the critical force is *∼*86 pN [41].

The nuclear envelope exerts equal and opposite radially inward forces on MT bundle ends, which produce a net force that moves the bundle center of mass toward the center of the nuclear envelope, but produces no net torque (Figure S1B). MT bundle minus ends interact with SPBs; this coupling to SPBs with their large drag breaks the symmetry between plus-and minus-ends and produces a net torque about the bundle centers. SPB drag on the minus ends of MT bundles tends to rotate the bundles away from alignment, leading to a torque

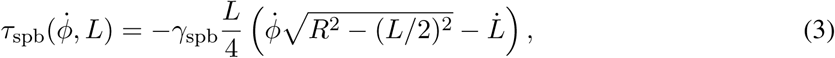

where *γ*_spb_ is the friction coefficient of SPBs, *R* is the radius of the nuclear envelope, *φ* is the bundle crossing angle, and 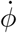 is the time derivative of *φ* (Figures 2A, S1A). The magnitude of the SPB torque *τ*_spb_ is proportional to *γ*_spb_, the average MT polymerization speed 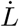, and bundle length *L* (Equation S13). However, longer MTs experience a greater parallel force (along the MT axis) from the nuclear envelope than perpendicular force (perpendicular to the MT axis), which slows MT polymerization, reduces SPB velocity, and therefore lowers the antialigning torque (Figure S1).

We calculate the average force and torque exerted by crosslinkers on MT bundles by considering the statistical mechanics of crosslinker binding, unbinding, and stretching/compression. Because crosslinker binding and unbinding kinetics are relatively fast (time scale of seconds) compared to spindle assembly (time scale of minutes), we make a quasi-steady-state assumption that crosslinker binding equilibrates given the instantaneous MT bundle positions. Therefore, we can determine crosslinker-induced forces and torques on MTs by computing the grand partition function and its derivatives. This approach is computationally inexpensive compared to calculation of single particle dynamics. Calculating forces and torques through the crosslinker partition function may be applied generally to filament networks with fast rearrangement of passive crosslinkers compared to the movement of filaments.

The single-crosslink partition function between two filaments is the integral of the Boltzmann factor 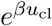, where *u*_cl_ is the crosslinker binding free energy, over all possible binding configurations [50, 51]

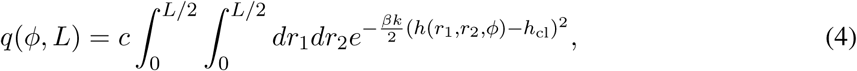

where *c* is the crosslinker concentration, the integrals over crosslinker endpoints *r*_1_, *r*_2_ extend over half the filament length from 0 to *L/*2, *k* is the crosslinker spring constant, *φ* is the angle between the two filaments, 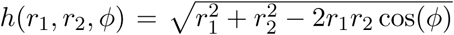 is the crosslinker length when the endpoints are at *r*_1_ and *r*_2_ along the two filaments, *h*_cl_ is the crosslinker rest length (at which it is neither stretched nor compressed), and *β* = 1*/*(*k*_B_*T*) is the inverse temperature (Figure 2B, Equation S17). We assume that crosslinkers do not interact with each other, which motivates the use of the grand canonical ensemble of a non-interacting gas with partition function

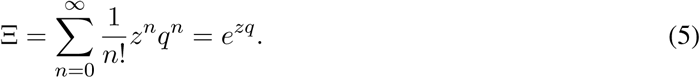

Here *n* is the number of bound crosslinkers and *z* is the fugacity 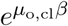, where *µ*_o,cl_ is the chemical potential of crosslinkers binding to the filaments. We can then determine crosslinker force and torque from derivatives of the grand potential

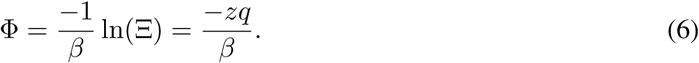

For example, the torque from all crosslinkers on MT bundles is

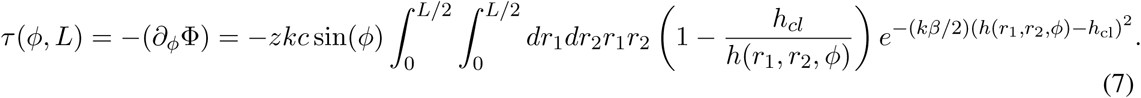

Torque due to crosslinkers can promote or oppose antiparallel alignment of MTs, because the magnitude and direction of the torque depends on MT length and the crossing angle (Figure 2). Crosslinkers can bind above, below, to the left, and to the right of the crossing point. Symmetry allows us to consider only the left/right and above/below cases: the angle in Equation 7 is *φ* for left/right and *π* - *φ* for above/below. As expected, a small angle between bundles produces greater crosslinker attachment, and therefore greater torque. Left/right binding typically produces aligning torque, although there can be exceptions when the crosslinkers are too compressed upon binding.

By using the Langevin equations for translational and rotational motion of filaments we can derive a system of integro-differential equations for *φ* and *L* (Section S1, Equations S21, S22)

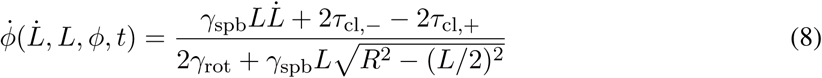

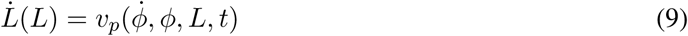

where *γ*_rot_(*L*) is the rotational drag coefficient of a filament of length *L* [89]. These equations can be solved numerically given initial *φ*_0_ and *d*_0_ (Figure 2A, Equation S2). The time evolution predicts the end configuration, either bipolar or aberrant (Figure 2C), (Supporting Information, section S1.1).

## Results and Discussion

### Phase diagram of spindle assembly

To compute a phase diagram for spindle assembly as a function of initial conditions, we numerically integrated Equations 8 and 9 (Supplementary Material). Dynamics that reach a bipolar final state have a decreasing crossing angle *φ* at the end of the integration, *dφ/dt <* 0 (Figures 2C, S2, S3, S4). Aberrant states occur for larger initial crossing angle because the aligning torque from crosslinkers is not able to overcome the anti-aligning torque from SPB drag (Figure 2D). For larger SPB separation, longer MTs provide more sites for crosslinker binding, while steric forces from the nuclear envelope are more parallel to the MT axis, slowing polymerization and reducing SPB drag. For MTs that span the nucleus, bipolarity occurs whenever *φ*_*o*_ *< π/*2, since the crosslinker aligning torque dominates. In fission-yeast spindle assembly, bundles tend to crosslink initially at short SPB separation [90]. Our results show that model parameters that favor bipolarity for small SPB separation will favor spindle assembly for a range of initial conditions. This result is sensitive to the MT polymerization rate: slower MT polymerization decreases the torque from SPBs, increasing the range over which spindles can reach bipolarity (Figure S5).

### Comparison to kinetic Monte Carlo-Brownian dynamics simulations

Our kinetic Monte Carlo-Brownian dynamics (kMC-BD) simulation model includes three-dimensional geometry, multiple sterically interacting MTs, and stochastic effects [50, 51], making it a useful point of comparison with the minimal torque-balance model (Figure 3, Movie S1). In contrast to the torque-balance model, kMC-BD simulations with the same parameters do not always end in the same state because of stochastic effects, including thermal motion, randomly generated MT nucleation sites and initial crosslinker location, and binding kinetics. These stochastic effects also allow spindles to escape aberrant states and become bipolar (Figure 3B). The three-dimensional geometry adds another degree of freedom, increasing the number of ways to escape from kinetic traps.

**Figure 3:**
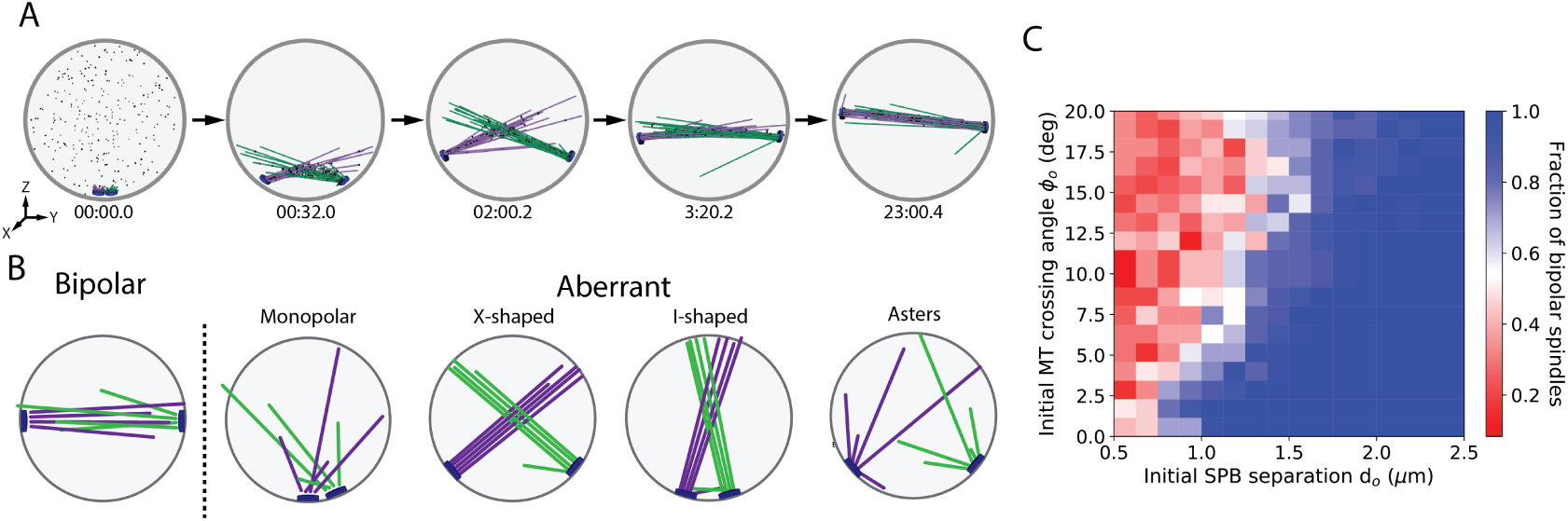
Kinetic Monte Carlo-Brownian dynamics simulation snapshots, typical final states of simulations, and phase diagram. A, Simulation snapshots of bipolar spindle assembly from an initial condition with adjacent SPBs. Times shown are in minutes, seconds, and tenths of seconds. B, Schematics of final states of simulations. C, Phase diagram of spindle formation failure: red indicates an aberrant end-state, blue the formation of a bipolar structure. Bipolar spindle fraction determined from 24 simulations at each data point. See Movie S1.

In our kMC-BD model, the total crosslinker number is fixed, and we compute the location of individual molecules, which allows crosslinker forces to fluctuate. Therefore, the overall force that crosslinkers can exert is constrained by the total crosslinker number. Crosslinkers must diffuse or unbind and rebind to apply force to recently overlapped MTs, processes that are modulated by thermal fluctuations, force, and MT configuration. The kMC-BD model becomes more similar to the quasi-steady approximation used in the torque-balance model for high crosslinker turnover and/or high bound diffusion coefficient. Unlike in the torque-balance model, the distribution of crosslinkers can be out of equilibrium, since small random thermal movements of MTs and SPBs occur on short time scales compared to the redistribution of crosslinkers. Out of equilibrium, crosslinkers exert restoring forces that maintain the network configuration despite random forces acting on MTs and SPBs. The stochastic binding kinetics allow kMC-BD simulations to escape kinetic traps but, if rearrangement is too slow compared to thermal motion, nonequilibrium effects can increase the strength of kinetic traps and the frequency of aberrant states. We quantified spindle assembly frequency as the fraction of simulations that end in a stable bipolar state after 24 minutes of simulated real time [51]. A higher spindle assembly frequency occurs in the phase diagram of the kMC-BD model (Figure 3C) for parameter sets that give bipolar spindle assembly in the torque-balance model (Figure 2D), demonstrating that the torque-balance model contains the key ingredients that control bipolar spindle assembly in the full kMC-BD model. The differences between these predictions highlight the importance of stochastic effects and crosslinker rearrangements in spindle assembly.

### Lengthening crosslinkers inhibits bipolar spindle assembly by over-bundling parallel MTs

Crosslinkers have a characteristic length, but it is not well understood whether changes in this length might affect spindle assembly. We assume Ase1 is around 40-55 nm long, based on the length of the human homolog PRC1 (37 nm long [91]) and kinesin-5 crosslinking motors (53 nm long [92]). In our model, crosslinkers are springs, so that stretched crosslinkers pull MTs together, whereas compressed crosslinkers push them apart. Crosslinker length determines the distance from the MT crossing point at which crosslinkers prefer to bind, which affects the amount of splay in MT bundles.

Short crosslinkers inhibit spindle assembly because crosslinkers primarily bundle neighboring MTs, limiting interdigitation of antiparallel MTs (Figure 4, movie S2). MT bundles become tightly bound and are less likely to become interdigitated with MTs from the other SPB, which decreases the strength of the crosslinker aligning torque (Figure 4A, bottom left inset).

**Figure 4:**
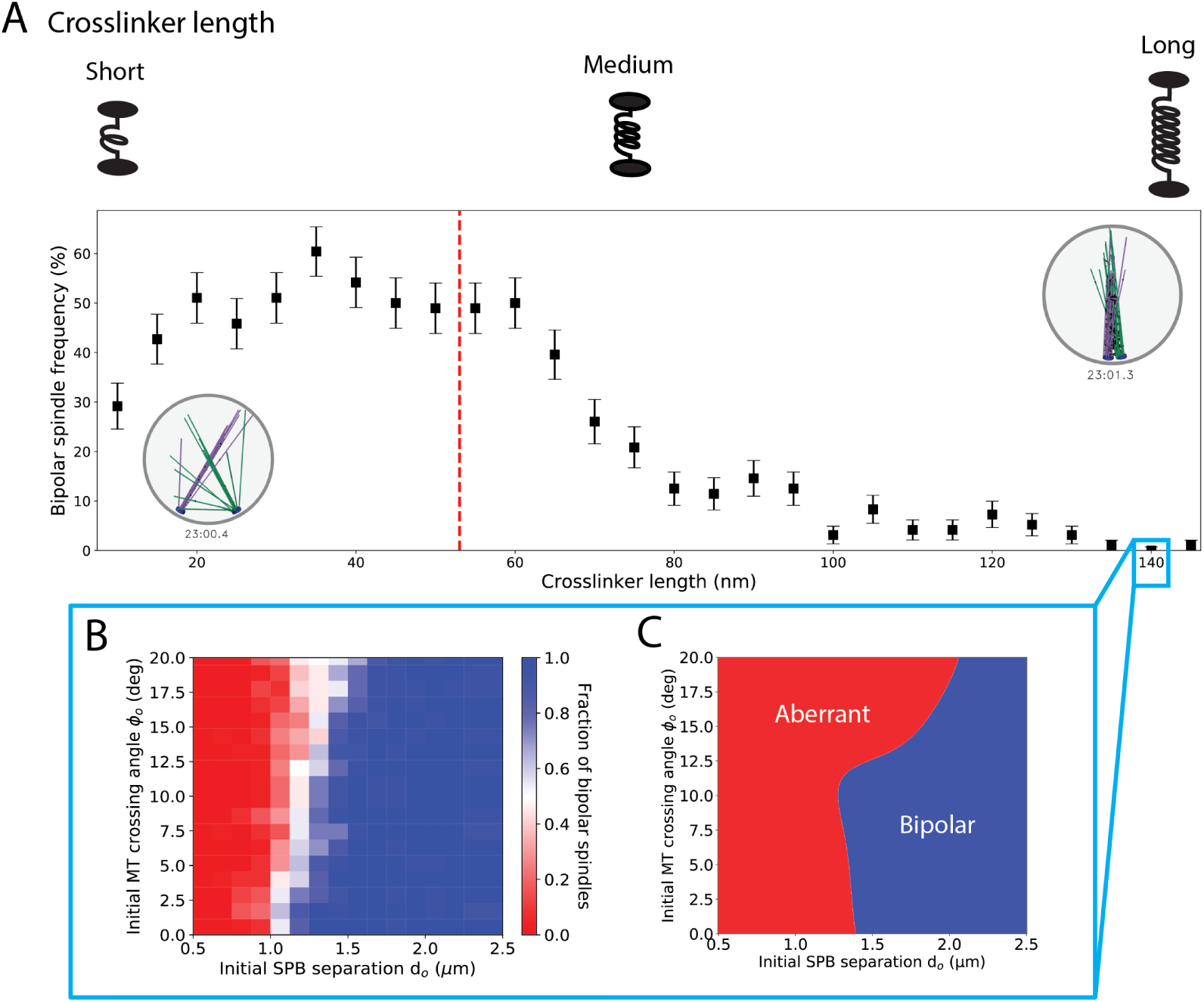
Crosslinker length of 20-60 nm is optimal for spindle assembly. A, Fraction of simulations that assemble a bipolar spindle as a function of crosslinker length. Top, schematic of crosslinker length effects. Inset, simulation snapshots of typical aberrant final states. Red dotted line indicates reference parameter value. B, C, Effects of long crosslinker length on spindle assembly, for crosslinkers of length 140 nm. B, Fraction of simulations that assemble a bipolar spindle as a function of initial SPB separation. C, Spindle assembly phase diagram from the torque-balance model with long crosslinkers. See Movie S2.

Increasing crosslinker length too far allows crosslinkers to interact with more MTs, which negatively impacts spindle assembly in two ways: crosslinking occurs when the SPB separation is small, and MT bundles form aberrant I-and X-shaped spindles (Figure 3B). Longer crosslinkers splay parallel bundles, limiting MT crosslinking near the dynamic ends of the bundles. Crosslinking between splayed bundles tends to trap the SPBs close together, leading to a narrow X-or I-shaped spindle (Figure 4A, insets, movie S2). These effects make I-and X-shaped spindles strong kinetic traps for long crosslinkers.

These results suggested that the defects of the model with long crosslinkers might be rescued by separating SPBs at the start of the simulation. To test this prediction, we began simulations with SPBs separated and allowed MTs and crosslinkers to interact while the SPBs were held in place for 1 second. Spindle assembly frequency increases sharply as initial SPB separation increases above around 1.2 *µ*m, similar to the location of the phase boundary in our torque-balance model (Figure 4B,C).

### Increased crosslinker turnover helps spindles escape kinetically trapped aberrant states

Because in previous work we found that crosslinker-only simulated spindles can become stuck in persistent monopolar states [51], we tested whether more rapid crosslinker turnover can promote bipolar spindle assembly by accelerating escape from aberrant states (Figure 5, Movie S3). Increasing turnover does not alter the average number of bound crosslinkers, but increases binding and unbinding (between one-and two-head bound states). With increased turnover, stretched or compressed crosslinkers more rapidly detach and can re-bind in states closer to mechanical equilibrium of the spring. Rapid turnover therefore makes random thermal forces more effective at repositioning MT bundles.

**Figure 5:**
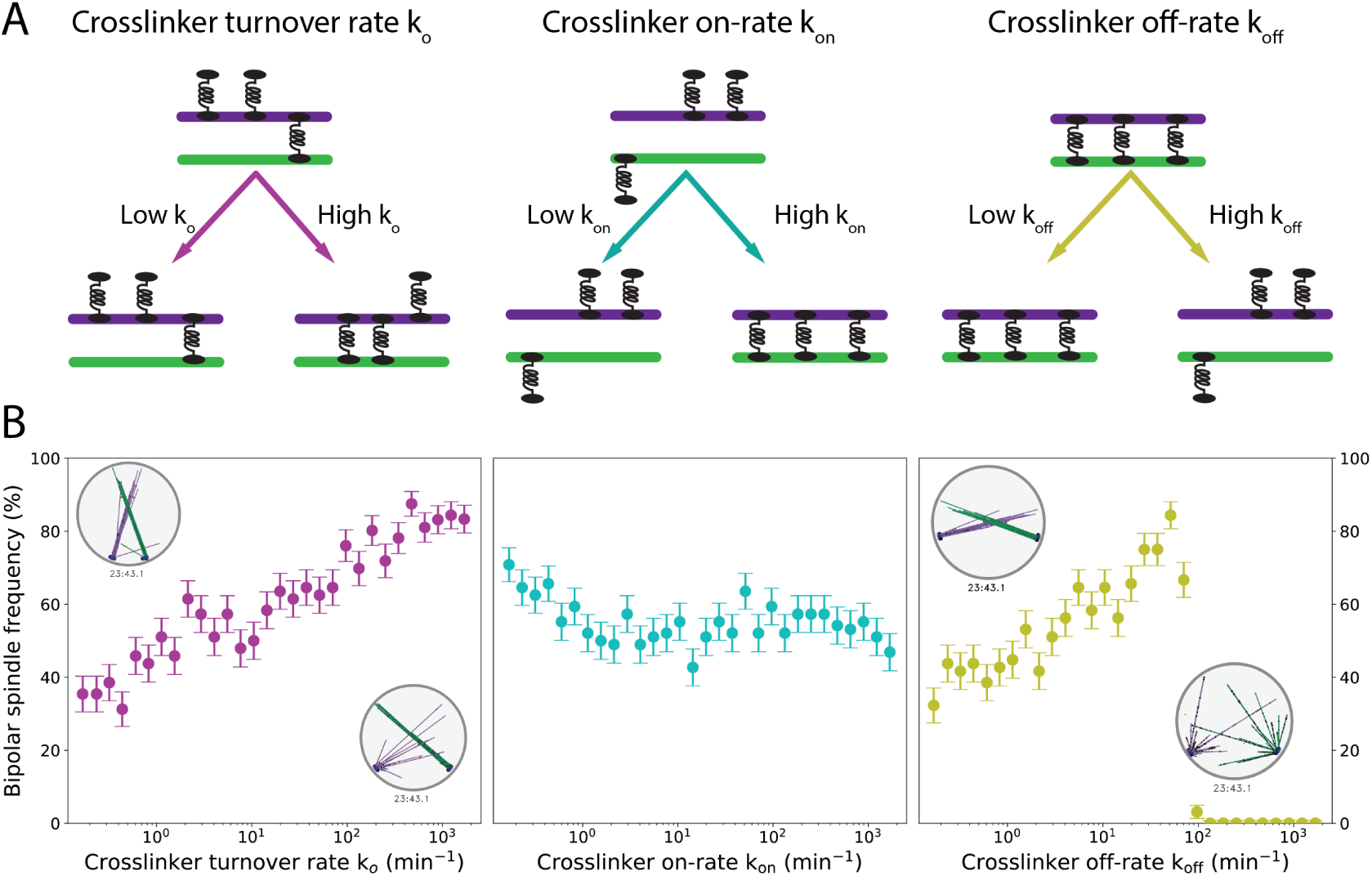
High crosslinker turnover facilitates spindle assembly. A, Schematic effects of varying crosslinker kinetic rates. Changing turnover (left) changes both on- and off-rates for transitions between one and two heads bound. Changing the on-rate (center) changes the binding rate from one head to two. Changing off rate (right) changes the unbinding rate from two heads to one. B, Fraction of simulations that assemble a bipolar spindle as a function of crosslinker kinetic rates. See Movie S3.

Similar to increasing the turnover rate, increasing the unbinding rate of doubly-bound crosslinkers facilitates the transition to a bipolar spindle by accelerating bundle reconfiguration. However, increasing the unbinding rate too much decreases the average number of crosslinks, leading to a critical value of unbinding rate above which the number of bound crosslinkers is too low to maintain bundle integrity (Figure 5B, Movie S3).

When crosslinker turnover is rapid, the kMC-BD simulation model more closely approximates the torque-balance model’s quasi-steady binding approximation. In this limit, crosslinkers rapidly change their configuration to the most statistically probable. In this case, we expect that fluctuations and aligning torque from crosslinkers will allow X-shaped spindles to form a bipolar spindle, given sufficient time. While not every simulated spindle reaches a bipolar state with high crosslinker turnover, the trends match this expectation.

### Crosslinker diffusion facilitates escape from kinetic traps by promoting crosslinker redistribution and MT reorientation

One-dimensional diffusion of bound crosslinkers repositions crosslinker heads on the MTs, which suggest that a higher diffusion coefficient might promote escape from aberrant states. Since diffusion is biased by force, when random thermal forces reorient MTs and extend/compress the crosslinkers, diffusion favors crosslinker motion that reduces this force. Therefore, diffusion modulates MT structural rearrangement: slow diffusion tends to inhibit MT reorientation, while fast diffusion tends to promote it (Figure 6, Movie S4).

**Figure 6:**
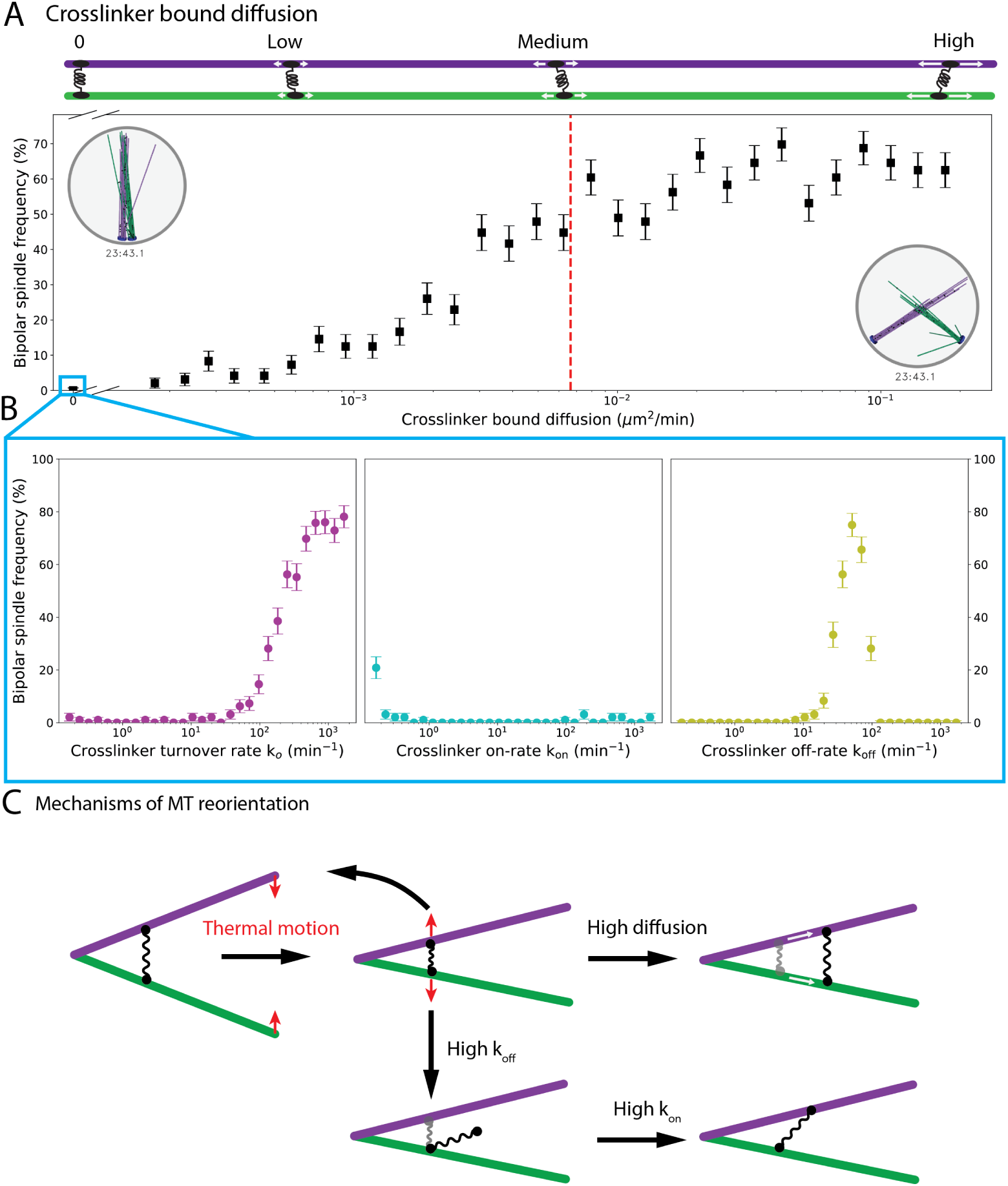
Bound crosslinker diffusion facilitates crosslinker redistribution and spindle assembly. A, Fraction of simulations that assemble a bipolar spindle as a function of bound crosslinker diffusion coefficient. Red dotted line indicates reference parameter value. Top, schematic of bound diffusion coefficient effects. Inset, simulation snapshots of typical aberrant final states. B, Fraction of simulations that assemble a bipolar spindle as a function of crosslinker kinetic rates, in the absence of bound crosslinker diffusion. Bipolar spindle formation is rescued by high crosslinker turnover. C, Schematic showing how high crosslinker turnover can rescue MT reorientation in the absence of bound diffusion. See Movie S4.

Increasing the doubly bound diffusion coefficient promotes spindle assembly, similar to the effects of increased turnover, but there are some differences. An increased diffusion coefficient by itself does not release crosslinked MTs, so does not lead to the separated asters seen for high crosslinker unbinding rate. In the opposite limit, when we completely remove doubly-bound crosslinker diffusion, bipolar spindles do not form, because crosslinkers do not rearrange quickly enough. Instead, these simulations produce long-lived I-or X-shaped spindles. In I-shaped spindles, the crosslinked bundles remain close together, since crosslinkers cannot migrate when SPBs fluctuate apart.

The similar increases in spindle assembly frequency for increasing turnover and diffusion suggest that changing turnover might be able to rescue the absence of diffusion: if binding kinetics are sufficiently rapid, one head can unbind and reattach in a different position, redistributing the crosslinkers. Consistent with this, high turnover rescues spindle assembly in simulations lacking bound diffusion (Figure 6B). Similarly, high crosslinker unbinding rate can rescue spindle bipolarity for a narrow range of values before reaching the critical value at which too few crosslinkers remain bound.

### Bipolar spindles form most readily when the parallel-antiparallel binding ratio is low but non-zero

Crosslinkers of the Ase1/PRC1/MAP65 family have a binding preference for antiparallel MT overlaps [14, 15, 93], but it has not been determined whether this bias affects crosslinker-mediated bipolar spindle assembly. Previous work has found that crosslinkers in this family favor antiparallel MT binding over parallel by a factor of two to ten [13]. In our model, we implement this effect as a binding enhancement that changes when the angle between MT axes is greater or less than 90 degrees. We then examined whether varying this binding preference affects spindle assembly (Figure 7).

**Figure 7:**
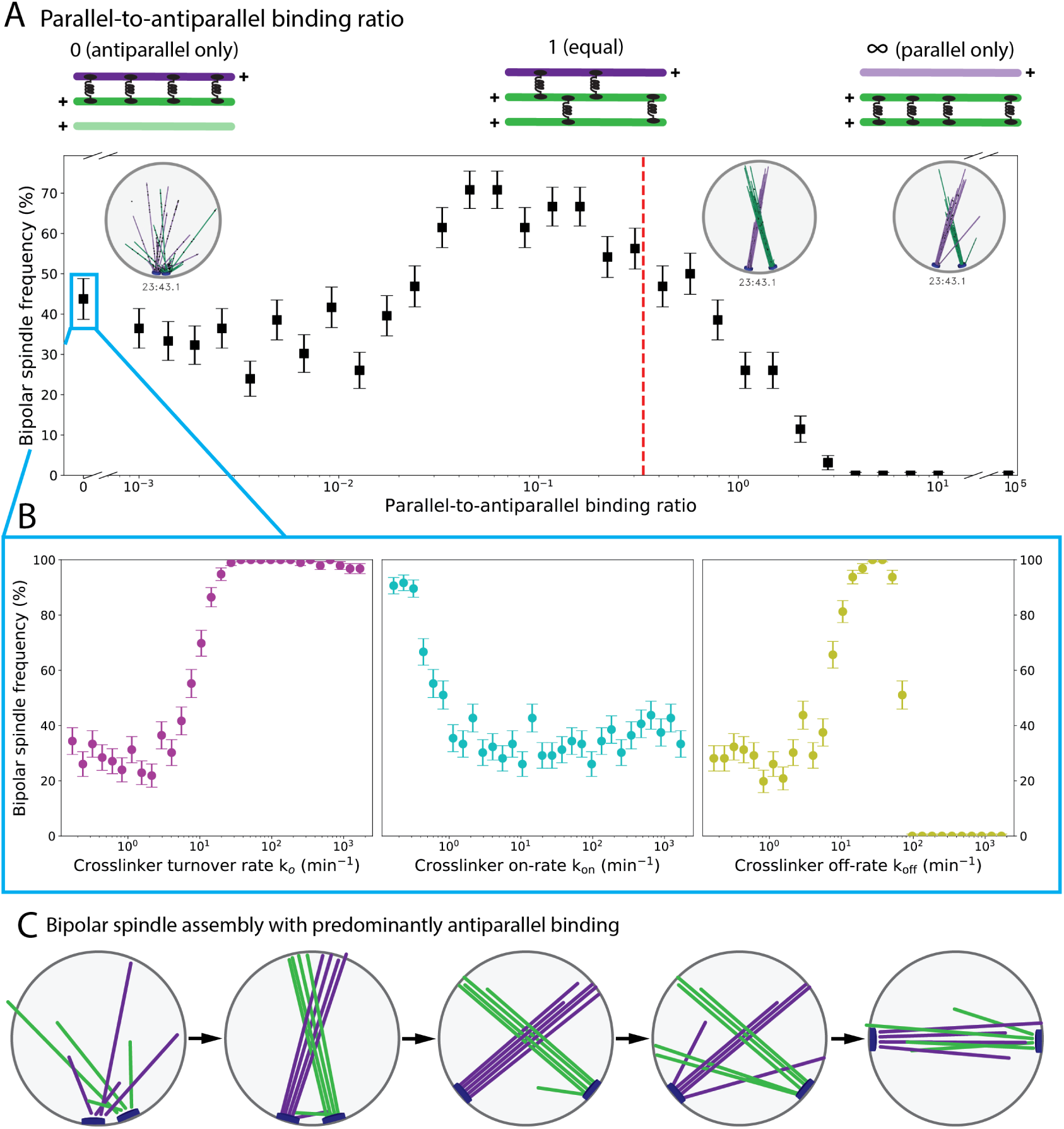
The crosslinker binding preference for antiparallel MTs promotes spindle assembly. A, Fraction of simulations that assemble a bipolar spindle as a function of the parallel-to-antiparallel binding ratio *α*. The red dotted line indicates the reference parameter value. Top, schematic of binding ratio effects. Inset, simulation snapshots of typical aberrant final states. B, Fraction of simulations that assemble a bipolar spindle as a function of crosslinker kinetic rates in the absence of parallel binding. C, Schematic of spindle assembly pathway for the optimal binding ratio. MTs crosslink parallel bundles, SPBs separate, then bundles interdigitate at shallower angle to form a bipolar spindle. See Movie S5.

For low parallel-antiparallel binding ratio (*α*=0–0.01), spindles can form because crosslinkers inhibited in parallel bundling avoid the X-shaped spindle. In this limit, failure of bipolar spindle assembly occurs sometimes because antiparallel bundling for very short MTs from adjacent SPBs leads to trapped monopolar spindles (Figure 7, Movie S5). Parallel crosslinking can promote spindle assembly by forming X-shaped spindles that allow the SPBs to separate, then interdigitate.

For low-intermediate parallel-antiparallel binding ratio (*α*=0.01–0.1), predominantly antiparallel binding with some parallel binding allows X-shaped spindles to transition to bipolar spindles, leading to the highest frequency of spindle assembly observed. Monopolar spindles tend to be short-lived, because parallel binding allows crosslinkers to migrate away from short antiparallel overlaps, promoting SPB separation (Figure 7C, Movie S5). Bundles then either break up to form a bipolar spindle, or rotate until the bundles are antiparallel.

For larger parallel-antiparallel binding ratio (*α*=0.1–0.3), the modest increase in parallel crosslinking favors the X-shaped spindle, inhibiting bipolarity. The lifetime of the X-shaped spindle increases with *α*. Similarly, when parallel and antiparallel binding are equally likely (*α*=1), X-shaped spindles typically form. However, random thermal forces can occasionally lower the angle between bundles, allowing antiparallel crosslinks and a bipolar spindle to form. Fully extended bundles at right angles to each other have, on average, balanced torques promoting alignment and anti-alignment. This balance can be broken if a thermal fluctuation rotates bundles toward antiparallel alignment.

For purely antiparallel binding (*α* = 0), increasing crosslinker turnover has a dramatic effect (Figure 7B). If turnover is sufficiently high, purely antiparallel-binding crosslinkers produce bipolar spindles for nearly every simulation. Varying turnover and unbinding rate show similar trends, except for the failure above the critical value for the unbinding rate at which most crosslinkers unbind. As for the other model perturbations discussed above, rapid rearrangement of crosslinkers allows spindles to escape kinetic traps and become bipolar.

Spindles typically transition from X-shaped to bipolar via two pathways: reconfiguration of crosslinkers, and aligning torques between bundled MTs (Figure 7C, Movie S5). Slow reconfiguration occurs when MT bundles move into antiparallel alignment due to thermal motion and crosslinker torque. The aligning torque pathway occurs more often when *α* is small, so that single polar MTs are common. When single MTs escape the main bundles, they can crosslink with MTs from the other SPB at a relatively shal-low angle. This allows the MTs to push against the nuclear envelope and separate SPBs until the main MT bundles also align into a fully bipolar spindle.

## Conclusion

Here we have developed a physical theory for cytoskeletal reorganization during fission-yeast mitotic spindle assembly (Figure 1) that incorporates the key ingredients of filament polymerization and depolymerization, crosslinking, steric interactions, and drag (Figure 2). Comparison of our minimal torque-balance model to full kMC-BD simulations that incorporate all stochastic effects (Figure 3, Movie S1) shows good agreement, demonstrating that the torque-balance model can illuminate the physical constraints on crosslinker-mediated spindle assembly. We studied specific individual perturbations to crosslinker length (Figure 4, Movie S2), binding kinetics (Figure 5, Movie S3), bound diffusion (Figure 6, Movie S4), and parallel versus antiparallel binding preference(Figure 7, Movie S5). The results demonstrate that crosslinkers that favor binding to antiparallel MT pairs and rapid redistribution of crosslinkers are crucial for bipolar spindle assembly from initially side-by-side SPBs.

In crosslinker-mediated spindle assembly, the crosslinker binding preference to antiparallel rather than parallel MT pairs is important for spindle assembly: bipolar spindles have more possible binding states between antiparallel MT pairs, so the antiparallel binding preference energetically favors bipolarity. However, this state must still be kinetically reachable from an initial condition in which spindle MTs are predominantly parallel. Therefore, crosslinker-mediated spindle assembly is vulnerable to kinetic traps at key stages of assembly (Figure 3). If antiparallel crosslinking predominates when SPBs are close to each other and crosslinker redistribution is too slow, spindles become trapped in a monopolar state. If parallel crosslinking predominates and crosslinkers either cannot bind to antiparallel MTs or redistribute too slowly, the frozen parallel bundles of X-or I-shaped spindles predominate. If too few crosslinkers are present, the spindle can fall apart into separated asters.

We further examined whether changing the binding and unbinding rates (Figure 5A, Figure 7B) individually has similar effects. Increasing the on-rate tends to lock in monopolar states, because it increases the total number of bound crosslinkers. For high binding rate, nearly all crosslinkers are bound to two MTs, preventing them from reorienting. By contrast, for low binding rate, crosslinkers can remain with one head bound to an MT while they diffuse, then reform a crosslink at another position. Our results therefore suggest that the rate of crosslinker rearrangement controls the rate of MT rearrangement, and thus the rate of bipolar spindle assembly.

Spindles avoid or escape kinetic traps when aligning torques overcome anti-aligning torques early in spindle assembly. The torque-balance model demonstrates that aligning torques are strongest for separated SPBs and low crossing angle between MTs. In some cases, aligning torques must overcome the anti-aligning torques early in our model simulations if the spindle is to become bipolar. Important stochastic effects that promote escape from kinetic traps include crosslinker redistribution and random thermal forces. Crosslinker redistribution is faster when crosslinker binding kinetics and/or bound diffusion are more rapid. Remarkably, increasing crosslinker turnover or diffusion can rescue defects in bipolar spindle assembly caused by crosslinkers of non-optimal length or exclusively antiparallel crosslinker binding.

The modeling techniques we use are generally applicable to cytoskeletal reorganization in which crosslinkers facilitate reorientation of filaments. This area is a frontier of cytoskeletal theory and model-ing, as the field confronts more challenging three-dimensional problems. Despite the complex dynamics and large-scale filament rearrangements that occur during bipolar spindle assembly, our work shows that the key physical effects can be understood both in detailed simulations and a minimal torque-balance model. The principles and modeling methods we describe are broadly applicable to cytoskeletal systems.

## Acknowledgements

The authors would like to thank Robert Blackwell for helpful conversations. This work was funded by NSF grants DMR1725065 (MDB), DMS1620003 (MAG and MDB), and DMR1420736 (MAG); NIH grants K25GM110486 (MDB), R01GM124371 (MDB); a fellowship provided by matching funds from the NIH/University of Colorado Biophysics Training Program (AL); and use of the Summit supercomputer, supported by NSF grants ACI1532235 and ACI153223.

## Author contributions

ARL, MAG, and MDB formulated the torque-balance model and code; ARL, CE, and MAG wrote the kMC-BD simulation code; ARL ran kMC-BD simulations; ARL, CE and MDB analyzed model results; ARL and MDB drafted the manuscript; ARL, CE, MAG, and MDB edited the manuscript.

## Competing interests

The authors declare that they have no competing interests.

## Data and materials availability

All data and computer code for this study are available on request from the authors.

