## Supplemental text and figures for "Theory of cytoskeletal reorganization during crosslinker-mediated mitotic spindle assembly"

### Supplemental material

March 1, 2019

#### S1 Torque-balance model

We derive a system of integro-differential equations for $\phi$ and $L$ by balancing the forces and torques applied to MT bundles (Figure S1). This model includes three forces: steric force between the nuclear envelope and MT ends, drag from the nucleoplasm and SPB translation, and crosslinker force between MT bundles. We label MT bundle length $L$ and MT crossing angle $\phi$ (Figure S1A). Then we find the normalized bundle length

$$l = \frac{L}{2R}, \tag{S1}$$

SPB separation

$$d(l, \phi) = 2R\cos\left(\cos^{-1}(l) + \frac{\phi}{2}\right), \tag{S2}$$

and angle between the SPB separation and SPB normal vectors

$$\theta(l, \phi) = \cos^{-1}(l) + \frac{\phi}{2}. \tag{S3}$$

Due to symmetry we need only consider the equations of motion of a single bundle: we assume that MTs overlap at their centers. This approximation is valid for large $d$ and small $\phi$ such that

$$\cos\left(\cos^{-1}\left(\frac{d}{2R}\right) - \frac{\phi}{2}\right) - \frac{d}{2R\cos(\phi/2)} \ll 1. \tag{S4}$$

Initial conditions are chosen so our simulations begin in this regime.

##### Force and torque balance

The first equation of motion for an MT bundle describes the force balance parallel to the bundle axis

$$0 = -\gamma_{\text{lin},\|} v_{\text{c},\|} - F_{+,\|} + F_{-,\|} + F_{\text{spb},\|}, \tag{S5}$$

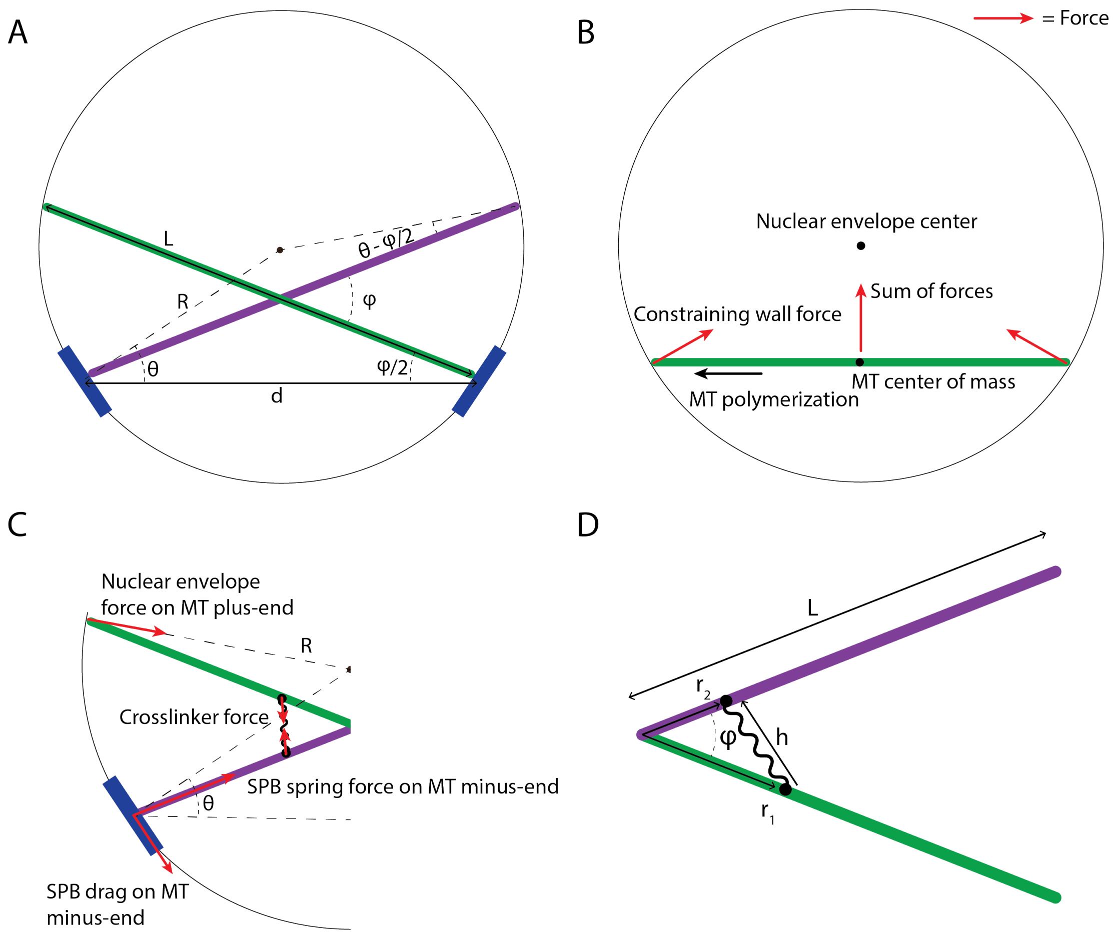


Figure S1: Schematic of the torque-balance model. A, Microtubule bundles, nuclear envelope, and spindle pole bodies. B, MT polymerization moves MT bundles toward the center of the nucleus. C, Forces exerted on crosslinked bundles. D, Crosslinked antiparallel microtubules.

where $F_{\pm,\parallel}$ is the parallel force component on the MT bundle ends due to the wall and $F_{\mathrm{spb},\parallel}$ is the parallel component of the SPB drag force (Figure S1B,C). The first term on the right hand side of Equation S5 is the drag force from the MT bundle moving parallel to its axis with $\gamma_{\mathrm{lin},\parallel}$ and $v_{\mathrm{c},\parallel}$ the parallel friction coefficient and bundle velocity. The bundle center of mass only moves parallel to its axis by polymerization; therefore, this drag force is approximately zero, i.e., $\gamma_{\mathrm{lin},\parallel} v_{\mathrm{c},\parallel} \approx 0$.

The second equation of motion balances the torque on the MT bundle

$$0 = -\tau_{\mathrm{drag,r}} + \tau_{+,\mathrm{wall}} - \tau_{-,\mathrm{wall}} + \tau_{\mathrm{spb}} + \tau_{\mathrm{cl},-} - \tau_{\mathrm{cl},+} \tag{S6}$$

where $\tau_{\mathrm{drag,r}}$ is the rotational drag from the nucleoplasm, $\tau_{\pm,\mathrm{wall}}$ is the torque from wall-end steric interaction, $\tau_{\mathrm{spb}}$ is the torque from the SPB on the bundle minus end, and $\tau_{\mathrm{cl},-}$ and $\tau_{\mathrm{cl},+}$ are the aligning and anti-aligning torque from crosslinkers. The nucleoplasm does not exert a torque since the bundle pivot point is at its center, the point of action of the total drag force, if Equation S4 holds.

##### Force

How MT bundles interact with the nuclear envelope determines the steric force on MT ends. MT growth speed depends on the parallel force exerted on its plus end. We model MT dynamics as in previous work, with polymerization speed $v_o$ and stall force $F_{\mathrm{s}}$ [87]

$$v_{\mathrm{p}}(l,\phi) = \frac{v_o}{\eta - 1}\left(\eta^{1-F_{+,\parallel}(l,\phi)/F_{\mathrm{s}}} - 1\right), \tag{S7}$$

where $\eta = e^{\delta F_{\mathrm{s}}/N k_b T}$, $\delta$ is the size of a tubulin dimer, 8 nm, $N$ the number of protofilaments in a microtubule, 13, $k_b$ the Boltzmann constant, and $T$ the temperature. Our model sets the nuclear envelope force to a constant directed toward the center of the nuclear envelope, $\vec{F}_+ = \text{const}\ \hat{r}$. The parallel component is

$$F_{+,\parallel} = F_+ l. \tag{S8}$$

As MTs grow, the angle between the nuclear envelope normal vector and the MT bundle axis decreases, which increases the parallel component of the wall force $F_{\pm,\parallel}$, and slows the polymerization and motion of the bundle (Figure S1B). We choose the wall force constant to be the stall force of MTs $F_+ = F_{\mathrm{s}}$, which prevents unbounded MT growth since at $l = 1, v = 0$.

The SPB drag force is determined by the SPB velocity on the nuclear envelope (Figure S1C)

$$F_{\mathrm{spb}} = \gamma_{\mathrm{spb}} R\dot{\theta}, \tag{S9}$$

where $\gamma_{\mathrm{spb}}$ is the friction coefficient [50]. We define $F_{\mathrm{spb}}$ so that decreasing $\theta$ produces a force with a negative component perpendicular to the MT bundle axis. With Equation S3 we re-write Equation S9 as

$$F_{\mathrm{spb}} = \gamma_{\mathrm{spb}} R\left(-\frac{\dot{l}}{\sqrt{1-l^2}} + \frac{\dot{\phi}}{2}\right), \tag{S10}$$

where the parallel component is

$$F_{\mathrm{spb},\parallel} = -\gamma_{\mathrm{spb}} R\left(-\frac{\dot{l}}{\sqrt{1-l^2}} + \frac{\dot{\phi}}{2}\right) l. \tag{S11}$$

This is negative because a decreasing $\phi$ (with no change in length) produces a positive parallel force along the MT axis.

The force on the MT bundle ends from the nuclear envelope is found using Equation S5 and $\gamma_{\mathrm{lin},\parallel} v_{\mathrm{c},\parallel} = 0$, so that

$$F_{-,\parallel} = F_{+,\parallel} - F_{\mathrm{spb},\parallel}. \tag{S12}$$

##### Steric and drag torque

Torque changes the orientation of the MT bundles and thus changes $\phi$. Positive torque increases $\phi$. We calculate torque, with the exclusion of that produced by crosslinkers, from the cross product of forces and the displacement of the point of action from the pivot. Since we approximate MT centers to be the pivot point, perpendicular components of the forces mentioned above multiplied by half the bundle length ($L/2$) give the magnitude of the torque. The torque from SPBs is

$$\tau_{\mathrm{spb}} = -F_{\mathrm{spb},\perp} L/2 = -\gamma_{\mathrm{spb}} l R^2 \left( \frac{\dot{\phi}}{2}\sqrt{1-l^2} - \dot{l} \right) \tag{S13}$$

whereas the torque applied to MT plus and minus ends is

$$\tau_+ = F_{+,\perp} L/2 = F_{+,\parallel} R\sqrt{1-l^2}, \tag{S14}$$

and

$$\tau_- = F_{-,\perp} L/2 = (F_{+,\parallel} - F_{\mathrm{spb},\parallel}) R\sqrt{1-l^2}. \tag{S15}$$

The rotational drag on MTs from the nucleoplasm is

$$\tau_{\mathrm{drag,rot}} = -\gamma_{\mathrm{rot}}\dot{\phi}, \tag{S16}$$

where $\gamma_{\mathrm{rot}}$ is the rotational friction coefficient for an MT bundle modeled as a spherocylinder of length $L$ [89]. This leaves only the torque from crosslinkers to completely define Equation S6.

##### Crosslinker torque and molecule number

The statistical properties of an ensemble of passive crosslinkers (the average number bound, torque, force, etc.) are calculated from the grand canonical potential (Equation 6) for indistinguishable crosslinkers binding to two filaments pivoting around a common origin (Figure S1D). We assume crosslinkers do not interact with each other and bind to both bundles simultaneously. For the geometry of Figure S1D, the partition function is

$$q = c\int_0^{L/2}\int_0^{L/2} dr_1 dr_2 e^{-\frac{\beta k}{2}(h(r_1,r_2,\phi)-h_{\mathrm{cl}})^2} \tag{S17}$$

where $c$ is the density of attachment sites and has the dimensions of inverse length squared, $k$ is the spring constant of a crosslinker, $h = \sqrt{r_1^2 + r_2^2 - 2r_1 r_2\cos(\phi)}$ is the crosslinker head separation, and $h_{\mathrm{cl}}$ is the equilibrium length of each crosslinker. The crosslinker binding affinity to two microtubules is the product of the fugacity $z$ (Equation 6) and $c$, which are never separated in this model.

The torque produced on one MT bundle is the negative derivative of the grand potential $\Phi$ (Equation 7). The average crosslinker number comes from the derivative of $\Phi$ with respect to the chemical potential $\mu$

$$\langle N\rangle = -\partial_\mu\Phi = (-\partial_\mu z)\partial_z\Phi = -\beta z\partial_z\Phi = zq. \tag{S18}$$

Crosslinkers also bind above and below the pivot point of Figure S1A. These crosslinkers produce the anti-aligning torque $\tau_{\text{cl},+} = \tau_{\text{cl}}(l, \pi - \phi)$ and contribute to the overall crosslinker number. Therefore, the total bound crosslinker number and torque are

$$\langle N_{\text{tot}} \rangle = 2\langle N_+ \rangle + 2\langle N_- \rangle, \tag{S19}$$

and

$$\tau_{\text{cl,tot}} = \tau_{\text{cl},-} - \tau_{\text{cl},+}. \tag{S20}$$

Equation S20 is then used in Equation S6 to complete the torque-balance model.

#### Equations of motion

Combining Equations S5, S6, and S7, we derive a system of equations for $\dot{\phi}$ and $\dot{l}$. By solving Equation S6 for $\dot{\phi}$ and plugging in the value $F_{-,||}$ from Equation S12 we arrive at

$$\dot{\phi}(\dot{l}, l, \phi, t) = \frac{2R^2\gamma_{\text{spb}} l\dot{l} + \tau_{\text{cl},-} - \tau_{\text{cl},+}}{\gamma_{\text{rot}} + R^2\gamma_{\text{spb}} l\sqrt{1 - l^2}}. \tag{S21}$$

Normalizing the MT polymerization speed from Equation S7 gives

$$\dot{l}(l) = \frac{v_o}{2R(\eta - 1)}\left(\eta^{1-l}\right). \tag{S22}$$

Equations S21 and S22 are numerically integrated in python using the `odeint` function from the `scipy.integrate` library. This code uses the Fortran `ODEPACK` library and LSODA integrator. The crosslinker torque from Equation 7 is computed by Gaussian quadrature using the `dblquad` method from `scipy.integrate`.

##### S1.1 Phase boundary determination

We classified spindles as being aberrant if the final $\dot{\phi}$ was positive and bipolar if negative (Figure S2, Figure S3, Figure S4). We computed a grid of binary results (aberrant/bipolar) for different initial conditions. To estimate the smooth phase boundary between aberrant and bipolar states, we used a Support Vector Clustering (SVC) algorithm with a Gaussian kernel from the `scikit-learn` python package. Initial condition parameter ranges were normalized to improve efficiency of SVC during the learning process and rescaled afterwards. The kernel was initialized with an amplitude of 1 and a variance of $\sigma^2 = 0.03125$.

### S2 Initialization of kMC-BD phase diagram simulations

Simulations that replicate the initial conditions of the torque-balance model start with SPBs set at separation $d_o$ with the MT minus ends anchored at random locations on the SPBs. MTs

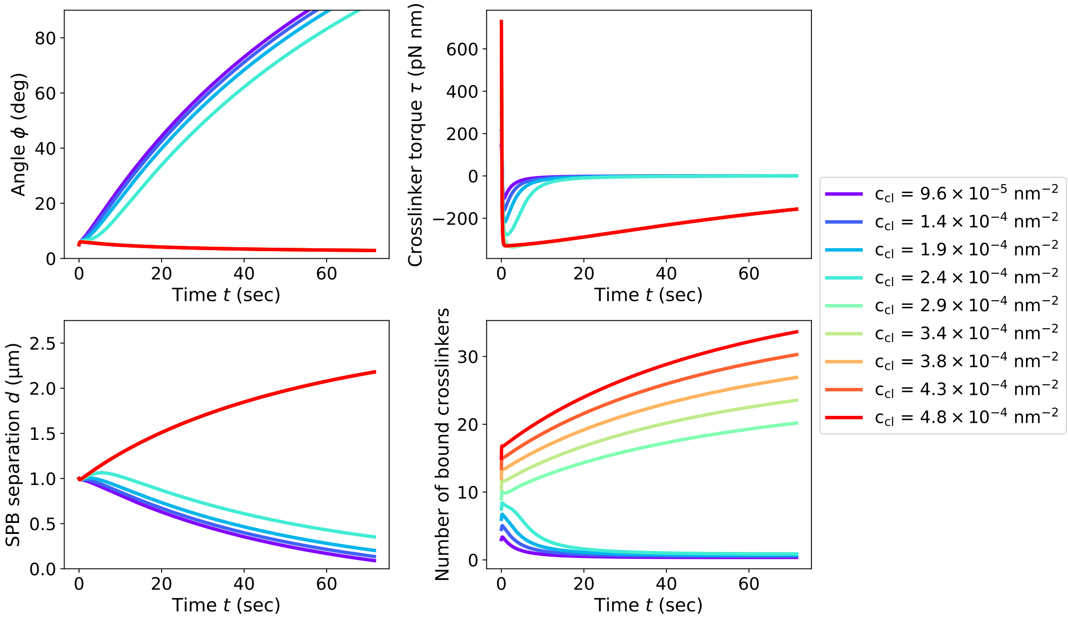


Figure S2: Time evolution of the torque-balance model with the parameters of Figure 2C, with the exception of the crosslinker binding affinity $c_{\text{cl}}$. The SPBs are initially separated by 1 $\mu$m and MTs cross at an angle of 5 degrees. Systems with $c_{\text{cl}} > 2.9 \times 10^{-4}$ nm$^{-2}$ assemble bipolar spindles because the crossing angle $\phi$ decreases while the SPB separation $d$ increases.

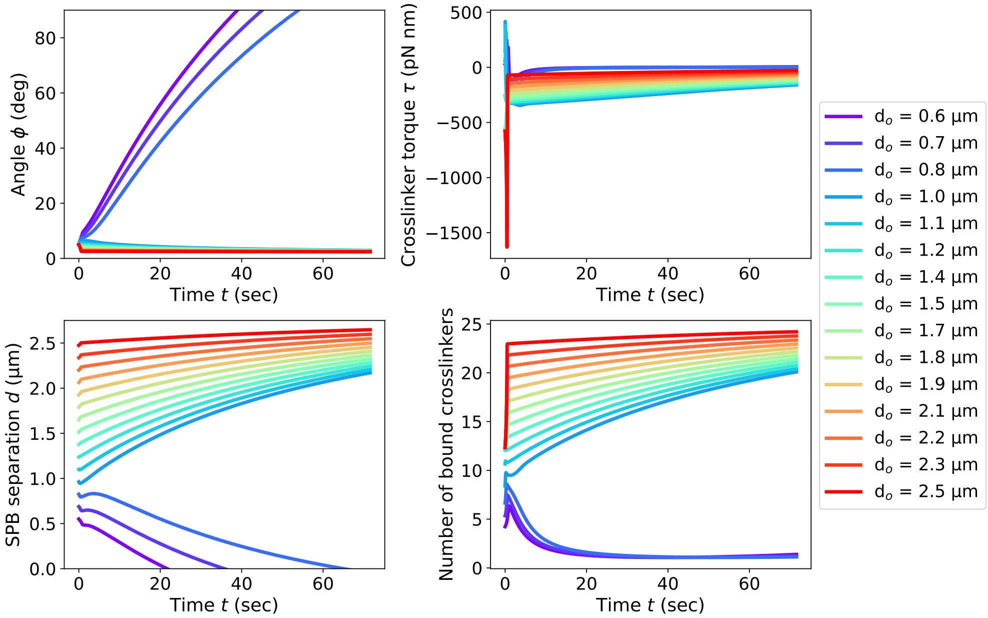


Figure S3: Time evolution of the torque-balance model with parameters of Figure 2C and MTs initially crossing at 5 degrees. The SPB separation is varied from 0.6 $\mu$m to 2.5 $\mu$m. The approximation that MTs bundle at their centers holds for systems of large SPB separation ($> 0.5$ $\mu$m) and small angle ($< 20$ degrees). Simulations that evolve to states outside this range give unphysical results at long time, as seen for $d_o < 1.0\ \mu m$. However, these simulations have monotonically increasing $\phi$, because aligning torque decreases with increasing $\phi$ after a certain $\phi$. This monotonicity of $\phi$ before becoming unphysical implies an aberrant final state for these simulations.

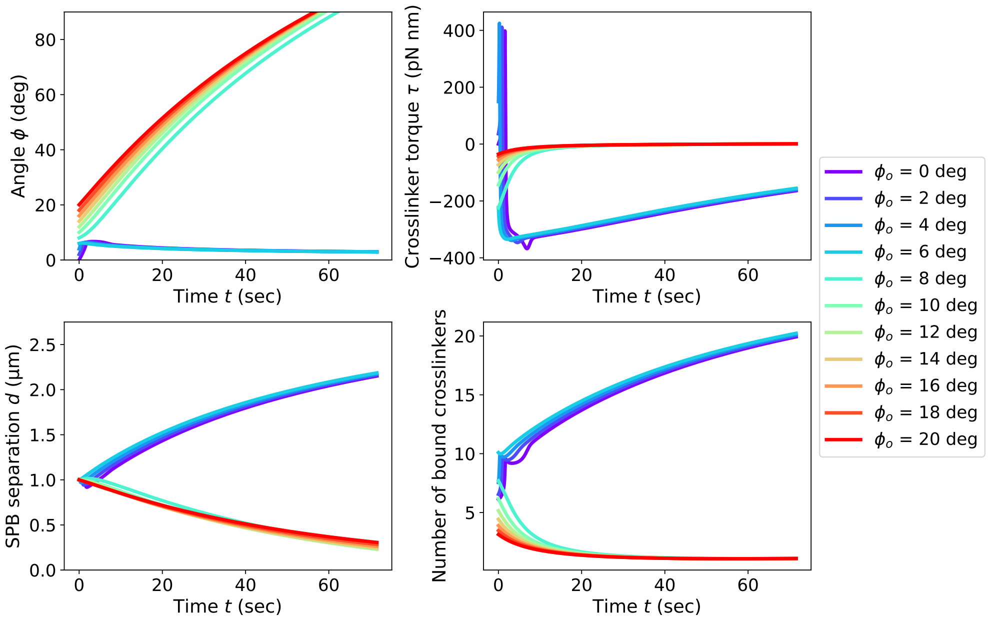


Figure S4: Time evolution of the torque-balance model with parameters of Figure 2C and SPBs initially separated by 1 $\mu$m. MTs initially cross at angles from 0 to 20 degrees. MTs crossing at angles greater than 8 degrees fail to form bipolar spindles. At small angle and short distance, crosslinkers exert an anti-aligning torque, since they are compressed. As bundle angle decreases, the crosslinking torque becomes positive, and the spindle becomes bipolar.

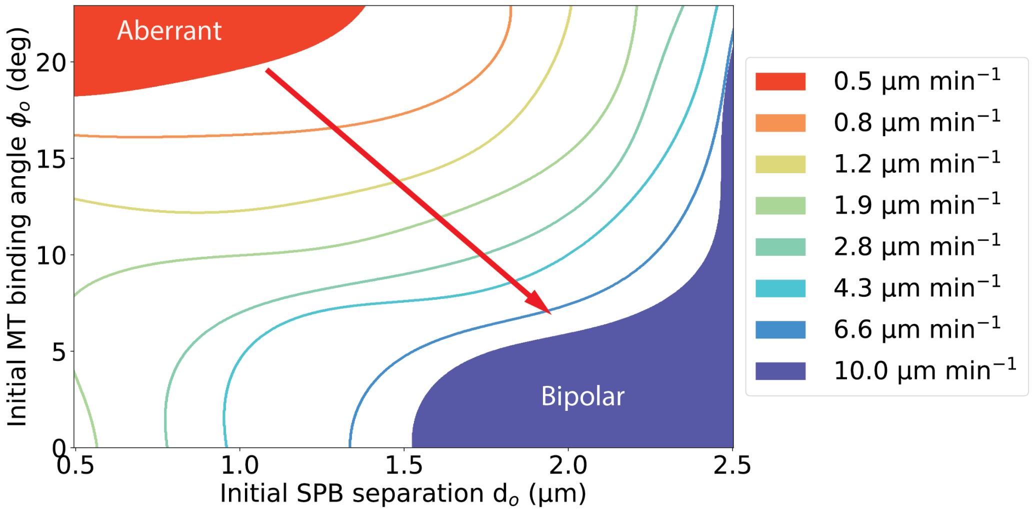


Figure S5: Change in spindle-assembly phase diagram in the torque-balance model as the average MT polymerization speed of microtubule bundles is varied. Lines represent the phase boundary between aberrant and bipolar spindles with bipolar spindles forming in lower right sections of the phase diagram. Filled regions label where aberrant spindles (red) or bipolar spindles (blue) for all parameters shown. Red arrow labels the expansion of the aberrant spindle region with increasing polymerization velocity.

| Parameter | Symbol | Value | Notes |
|---|---|---|---|
| Nuclear envelope radius | R | 1.375 $\mu$m | [75] |
| MT diameter | $\sigma_{MT}$ | 25 nm | [94] |
| MT angular diffusion coefficient | $D_\theta$ | Varies with MT length | [75] |
| Force-induced catastrophe constant | $\alpha_c$ | 0.5 pN$^{-1}$ | [87, 95] |
| Membrane tube radius | $f_{\text{tube}}$ | 87.7 nm | [96, 97] |
| Asymptotic wall force | $f_{\text{w}}$ | 2.5 pN | [96, 97] |
| **Spindle pole bodies** | | | |
| Diameter | $\sigma_{\text{SPB}}$ | 0.237 $\mu$m | [49] |
| Bridge size | – | 75 nm | [49] |
| Tether rest length | $R_0$ | 50 nm | [98, 99] |
| Tether spring constant | $K_0$ | 0.67 pN nm$^{-1}$ | Typical value of protein spring constant, cf. [50] |
| Number of MTs per SPB | $N_{MT}$ | 14 | [49] |
| Translational diffusion coefficient | $D_t$ | $4.5 \times 10^{-4}$ $\mu$m$^2$s$^{-1}$ | [50] |
| Rotational diffusion coefficient | $D_{\theta,\text{spb}}$ | 0.017s$^{-1}$ | [50] |
| **Dynamic instability** | | | |
| MT growth speed | $v_{g,0}$ | 4 $\mu$m min$^{-1}$ | [76] |
| MT shrinking speed | $v_{s,0}$ | 6.7 $\mu$m min$^{-1}$ | [76] |
| Catastrophe frequency | $f_{c,0}$ | 6.07 min$^{-1}$ | [76] |
| Rescue frequency | $f_{r,0}$ | 0.71 min$^{-1}$ | [76, 51] |
| Growth speed stabilization | $s_{vg}$ | 1.5 | Estimated based on model from [51] |
| Shrinking speed stabilization | $s_{vs}$ | 0.1 | Estimated based on model from [51] |
| Catastrophe frequency stabilization | $s_{fc}$ | 0.1 | Estimated based on model from [51] |
| Rescue frequency stabilization | $s_{fr}$ | 20 | Estimated based on model from [51] |
| Stabilization length | $s_\ell$ | 25 nm | Estimated based on model from [51] |
| Minimum MT length | $L_{\min}$ | 50 nm | Value chosen for numerical stability |
| MT stall force | $f_{\text{s}}$ | 14.6 pN | [87] |

Table S1: Microtubule, nuclear enevelope, and spindle pole body parameters used in kMC-BD model. MTs are modeled as rigid rods interacting with each other by a WCA potential. The plus-ends of MTs are directed radially inward by a non-monotonic force [50, 96] when they exceed the boundary of nuclear envelope. SPBs are constrained to the surface of the nuclear envelope on which they can diffuse. SPBs are attached to the minus-ends of MTs by a Hookian spring force.

are tilted away from the SPB normal vector so that the bundles cross at the desired $\phi_o$. All MT lengths are initialized so that an MT attached to an SPB center makes contact with the nuclear envelope. The simulation then initially runs for 1 second so that MTs do not overlap and crosslinkers bind to MTs. During the initialization, SPBs are held fixed and MTs remain at their initial length. Afterward, SPBs are released and MTs become dynamically unstable.

| **Parameter** | **Symbol** | **Value** | **Notes** |
|---|---|---|---|
| Available molecules | $N_{\rm tot}$ | 250 | [100] |
| One-dimensional effective concentration | $c_{c,2}$ | 0.4 $\rm nm^{-1}$ | [17] |
| Spring constant | $K_c$ | 0.2047 pN $\rm nm^{-1}$ | [17] |
| Diffusion constant (solution) | $D_{\rm free}$ | 4.5 $\mu\rm m^2$ $\rm s^{-1}$ | [101] |
| Singly bound diffusion constant | $D_{\rm sb}$ | 0.1 $\mu m^2$ $\rm s^{-1}$ | [17] |
| Doubly bound diffusion constant | $D_{\rm db}$ | 0.0.0067 $\mu m^2$ $\rm s^{-1}$ | Same as the singly bound hopping rate; [50] |
| Singly bound off-rate | $k_1$ | 0.1 $\rm s^{-1}$ | [102] |
| Doubly bound off-rate | $k_2$ | 0.05 $\rm s^{-1}$ | [17] |
| Parallel-to-antiparallel binding ratio | $\alpha$ | $1/3$ | [102] |
| Unbinding load sensitivity | $\lambda$ | 0.01626 | [50] |

Table S2: Passive crosslinker parameters used in kMC-BD model. Crosslinkers can be unbound, have one head bound, or have two heads bound.
